# Predictable induction responses of gut prophages

**DOI:** 10.64898/2026.06.25.734096

**Authors:** Laura Avellaneda-Franco, Sofia Dahlman, Jodee A. Gould, Denis Korneev, Remy B. Young, Emily L. Rutten, Samuel C. Forster, Jeremy J. Barr

**Author notes:** **Correspondence to**: Jeremy J. Barr. 25 Rainforest Walk, Clayton, Monash University, VIC Australia 3800. E.

## Abstract

Temperate bacteriophages are dominant members of the human gut microbiome that can infect and lyse their bacterial hosts or integrate as prophages. During this integrated state, prophages exhibit extensive control over host physiology and lysis via induction. Here, we studied a diverse collection of *Bacteroidales* isolates, which are amongst the most abundant bacterial orders within the human gut, identifying 902 high-quality prophage genomes present within 305 isolates, 240 of which were poly-lysogens. Despite their prevalence, our understanding of the function and induction triggers of prophages is limited. To predict prophage induction, we employed an iterative profile Hidden Markov Model search across divergent bacterial hosts to identify prophage regulatory components. We found 197 *Bacteroidales* prophages encoding complete CI-like repressor proteins, which initiate induction upon DNA damage. We selected *Bacteroides thetaiotaomicron* strain Bt_806 to characterise further as it harboured six diverse prophages, including the prevalent and abundant prophage LoVE, which was the only integrated prophage encoding a complete CI-like repressor. Transcriptomics revealed phage LoVE was routinely induced upon DNA damage, while the five co-habiting prophages remained stably integrated yet exhibited transcriptionally active genes associated with regulation, prophage maintenance, and uncharacterised functions. Finally, we selected an additional eleven *Bacteroidales* poly-lysogens, confirming that integrated prophages encoding complete CI-like repressors were reliably induced upon DNA damage. Together, we demonstrate that mechanistic understanding of prophage induction linked with identification of regulatory genes enables selective and predictable induction of gut prophage species as a potential tool to modulate the microbiome.

## Introduction

Bacteriophages, or phages for short, are the most abundant viruses in the human gut^1–3^. As viruses of bacteria, phages can infect and kill their host bacterial cells, releasing new viral progeny through the lytic cycle^4^. Temperate phages can delay replication via the lytic cycle by entering into lysogeny, whereby they establish as an integrative prophage element within their bacterial host chromosome^4,5^. Lysogeny is predicted to be the predominant phage lifecycle within the healthy adult human gut. In the human gut, prophage induction, which refers to the event by which prophages re-enter the lytic cycle typically in response to stress and environmental cues^6^, is estimated to occur at low yet consistent rates of ∼0.001-0.01 induction events per bacterium per day^7^. Thus, while gut bacterial cells outnumber free virions by ∼100-fold^7^, the majority of gut bacteria are lysogens^8^ (i.e. carrying prophages) and often poly-lysogens, which further contributes to the higher ratio of phage-to-bacterial genomes within this environment^7,9^. Higher relative abundance of temperate phage virions in the human gut has been observed during specific stages of life, including during infancy^10,11^ and in centenarians^12^, and has been associated with disease states, such as inflammatory bowel disease^13^. These associations implicate temperate phages, their induction, and putative functions with compositional and functional alterations of the gut ecosystem^14,15^. Understanding prophage induction triggers, their genetic signatures, and the molecular mechanisms that regulate it could prove impactful knowledge for the development and maintenance of a healthy human gut microbiome^16^.

*Bacteroidales* species are amongst the most abundant bacterial orders within the human gut^17,18^. Among *Bacteroidales*, *Bacteroidaceae* lineages have coevolved with hominids across hundreds of thousands of host generations, establishing enduring colonization within the human gut and fuelling metabolic networks that shape a web of microbial interactions^19,20^. Prophages are widespread within gut *Bacteroidales* isolates, with a recent study finding >90% of *Bacteroidota* isolates were lysogens and >70% of said isolates were poly-lysogens^8^. Despite the prevalence of *Bacteroidales* prophages within the gut, a mechanistic understanding of how they influence bacterial host physiology and switch from lysogenic to lytic cycles remains a significant knowledge gap.

Prophage induction genetics are well established in the model *Escherichia coli* phage Lambda. Here, the genetic switch is a complex gene circuit that exhibits regulatory control over lytic and lysogenic cycles^21–23^. Within it, the CI repressor protein plays two central roles^22^, one as the repressor of lytic cycle genes, and the other as a self-inactivator^24^. Each of these roles is separately accomplished by two different structural domains within the CI repressor protein; the N-terminal DNA-binding domain (NTD), which acts as the repressor, and the C-terminal catalytic domain (CTD), which is a peptidase that functions as the self-inactivator^25^ (**Fig. 1A**). During stable lysogeny, the NTD repressor binds to the prophage operator of lytic genes to repress their expression^25,26^. For induction, the CTD peptidase is activated in response to RecA-DNA filaments, which accumulate following host DNA damage, leading to the autocleavage of the repressor^25,26^. As a result, the induction of prophage Lambda first requires DNA damage-dependent activation of the CTD self-inactivator domain, which cleaves the CI protein, leading to the transcriptional activation of lytic genes and re-entry into the lytic cycle via induction. As has been proposed for other taxa^27^, here, we hypothesised that *Bacteroidales* prophages, despite their evolutionary divergence, contain core components of the Lambda-like genetic switch, which can be used to predict prophage induction in the gut.

**Figure 1.**
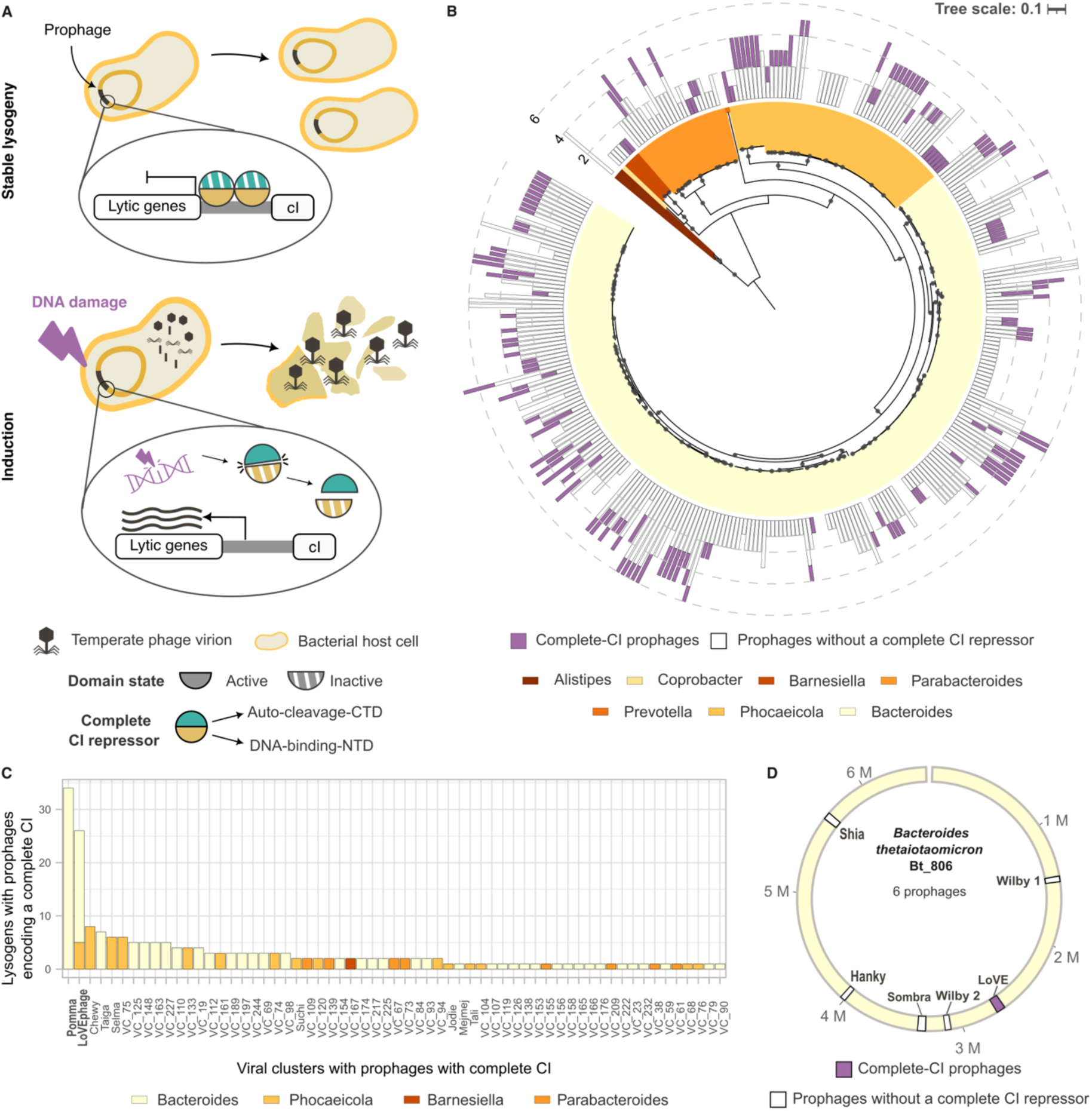
*Bacteroidales* prophages encode Lambda CI-like repressors. **A)** Schematic representation of prophage Lambda encoding the complete CI repressor and its function during stable lysogeny and induction following DNA damage. **B)** A phylogenetic tree of 335 *Bacteroidales* isolates from the human gut built on a 120-core bacterial gene alignment generated by GTDB-Tk^35^. Internal black dots indicate bootstrap values greater than 95% (using the UFBoot2 algorithm^36^). Each isolate is coloured based on its genus. The white bars show the number of prophages per isolate without a complete CI-like repressor, which includes prophages that only encode the NTD of the CI-like repressor or prophages that do not encode a CI-like repressor at all. The purple bars represent the number of complete-CI prophages per isolate that encode both functional domains. **C)** The prevalence of viral clusters (VCs) that encode a complete CI-like repressor is indicated by the number of lysogens in which each VC is present. **D)** Schematic representation of the Bt_806 genome and its six prophages, with phage LoVE being the only one to encode a complete CI-like repressor.

The present work utilized a diverse collection of 335 human gut *Bacteroidales* isolates to characterise the biology and induction of their prophages. We confirmed that *Bacteroidales* prophages encoding functionally complete CI-like repressors (i.e., containing both NTD and CTD) were prevalent in our collection. Next, we studied *Bacteroides thetaiotaomicron* strain Bt_806, a gut poly-lysogen carrying six complete prophages, to test whether the presence of a complete CI-like repressor was a reliable indicator of prophage induction upon DNA damage. Following the addition of the DNA-damaging agent Mitomycin C (MMC), we observed transcriptional activation of prophage LoVE, which was the only integrated prophage in isolate Bt_806 encoding a complete CI-like repressor. The remaining five prophages were not induced upon addition of MMC, yet exhibit transcriptionally active genes during stable lysogeny, indicative of transcriptional interaction with the host cell. To increase the robustness of this study, we selected an additional eleven *Bacteroidales* poly-lysogens from the gut, confirming that prophages encoding complete CI-like repressors were induced upon DNA damage. In summary, we provide evidence linking the identification of phage regulatory genes with predictable control over induction. We propose that this concept can be extended to other phage-host pairs in the gut to enable selective induction of phage species as a potential microbiome manipulation tool.

## Results

### A diverse collection of human gut *Bacteroidales* prophages encodes complete CI-like repressors

*Bacteroidales* prophages are prevalent within the human gut microbiome; however, the molecular mechanisms behind their induction remain poorly understood. Here, we employed 335 human gut *Bacteroidales* isolates (3 *Alistipes*, 239 *Bacteroides*, 5 *Barnesiella*, 1 *Coprobacter*, 26 *Parabacteroides*, 60 *Phocaeicola*, and 1 *Prevotella*; **Fig. 1B** and Supplementary Table 1) from the Australian Microbiome Collection Culture (AusMiCC) to predict prophage regions and identify their repressors, which function as regulators of induction. Combining manual curation with Virsorter2^28^, CheckV^29^, and DRAM-v^30^, we predicted 902 prophage regions and their respective proteome. High-quality, complete, predicted prophages were found within 91% of the *Bacteroidales* isolates (305/335), with 240 of the isolates being poly-lysogens (i.e. carrying multiple prophages). The 902 prophage regions were grouped into 248 viral clusters, with each cluster sharing 95% average nucleotide identity (ANI) over 85% aligned fraction (AF)^31^. Induction of 73 of the predicted prophages was experimentally confirmed in a prior study^8^. These 73 inducible prophages belonged to 22 viral clusters (VCs), which comprised a total of 310 predicted prophages, suggesting that at least ∼34% of the predicted prophages (310 out of 902), or ∼9% of the VCs within our collection (22 out of 248), were inducible.

Next, we sought to identify CI-like repressors within *Bacteroidales* prophages. We hypothesized that these regulatory domains shared conserved amino acid residues with the well-characterised CI repressor from the *E. coli* phage Lambda, despite divergence in their sequences. We employed an iterative profile Hidden Markov Model (profile-HMM) search method, JackHMMER^32^, to capture *Bacteroidales* proteins that share conserved residues with phage Lambda’s CI repressor protein (UniProt ID: P03034). This initial search yielded zero matches, suggesting evolutionary divergence among *Bacteroidales* and Lambda CI-repressors. To increase the phylogenetic resolution and, with it, improve the profile-HMM sensitivity, we included 654 AusMiCC gut bacterial isolates from four additional phyla (127 *Actinomycetota*, 353 *Bacillota*, 163 *Pseudomonadota*, and 11 *Fusobacteriota*, Supplementary Table 2) into the analyses. After predicting 1,271 prophage-regions within 543 (83%) of these additional gut isolates, the proteome of all predicted prophages was employed to build the profile-HMM using seven rounds of iterations (see methods). This extended search found 413 gut prophages across five phyla that encoded complete CI-like repressor proteins (9 *Actinomycetota*, 63 *Bacillota*, 142 *Pseudomonadota*, 2 *Fusobacteriota*, and 197 *Bacteroidota;* Supplementary Table 3 and Supplementary Table 4), which contained both the NTD repressor and CTD self-inactivator domains. Additionally, we found 1,104 gut prophages with CI-like repressors only encoding the NTD repressor domains (71 *Actinomycetota*, 448 *Bacillota*, 143 *Pseudomonadota*, 13 *Fusobacteriota*, and 429 *Bacteroidota;* Supplementary Table 3 and Supplementary Table 4), suggesting these phages encode novel or unidentified activators of induction.

To examine evolutionary relationships between prophage-encoded complete CI-like repressors, we constructed a phylogenetic tree of all identified repressor proteins (Supplementary Fig. 1A). The phylogenetic tree showed potential divergence in the evolutionary paths due to the weak support for the deep nodes, but nevertheless, recent relationships were resolved with higher confidence. Focusing on *Bacteroidales*, we found 197 prophages encoding complete CI-like repressors (complete-CI prophages), containing both NTD repressor and CTD self-inactivator domains. The complete CI-like repressors encoded by *Bacteroidales* prophages were typically located at the ends of the prophage-genome (Supplementary Fig. 1B) and formed a clade (conservative bootstrap 84) that was distinct from the classical phage Lambda CI protein. Taken together, these results indicate that ∼22% of *Bacteroidales* prophages (197 out of 902) encoded core components of the phage Lambda-like genetic switch at the ends of their genomes, which is suggestive of their inducibility upon DNA damage.

To select a gut prophage to study the association between repressor proteins and prophage induction, we assessed the prevalence of complete-CI prophages within the AusMiCC isolate collection. We found that the 197 *Bacteroidales* complete-CI prophages, which were grouped into 61 VCs, were found within roughly half of the *Bacteroidales* lysogens within this study (154/305, purple bar **Fig. 1B**). We observed that 59 out of the 61 VCs containing complete-CI prophages were present in less than ten lysogens within our collection, with only two of the 61 VCs being present in more than 25 *Bacteroidales* lysogens (**Fig. 1C**). These two prevalent VCs comprised members of the previously characterised phage Pomma and LoVE VCs, members of which had previously been confirmed as inducible in response to known DNA damaging agents (i.e., mitomycin C and Hydrogen peroxide)^8^. LoVE prophages belong to the proposed “*Quimbyviridae*” family and are common in the human gut, infecting hosts from the *Phocaeicola* and *Bacteroides* genera^2,8,33^. All LoVE prophages within our collection encoding complete CI-like repressors were found within poly-lysogenic *Bacteroidales* isolates. Thus, we selected prophage LoVE as an experimental model to test whether bioinformatic identification of a complete CI-like repressor is a reliable predictor of DNA damage-driven prophage induction within gut bacterial hosts.

### *Bacteroides thetaiotaomicron* Bt_806 and prophage LoVE as an experimental model to study prophage induction

We identified prophage LoVE as a prominent viral cluster that encodes a complete CI-like repressor, which we hypothesize is a predictor of DNA damage-driven prophage induction within *Bacteroidales*. To test this, we selected the poly-lysogen *Bacteroides thetaiotaomicron* Bt_806 (AusMiCC ID: CC00806), which had a genome size of 6.5 M bp and contained two plasmids and six integrated prophages (**Fig. 1D**). These six integrated prophages belong to five different VCs: prophage LoVE, which was the only integrated prophage to encode a complete CI-like repressor, prophage Hanky from Hankyvirus VC that are amongst the most prevalent *Bacteroidales* prophages in the gut^2,33,34^, two prophages from the recently described Sombra and Shia VCs^8^, and finally two Wilby prophages that were >99% genetically identical. All Bt_806 prophage genomes were annotated and predicted to be complete (98.7% mean completeness with a standard deviation of 1.34) (Supplementary Fig. 2). Furthermore, induction of prophages belonging to these five VCs has been previously validated in other isolates^8^, suggesting that all Bt_806 prophages were functional and can undergo lytic replication. We also note that the Bt_806 host genome contained an additional complete CI-like repressor that was not associated with any of the six integrated prophages, along with 22 antiphage defence systems. In summary, isolate Bt_806 harbors six complete prophages, with only phage LoVE encoding a complete CI-like repressor protein. To note, the remaining Bt_806 prophages encode incomplete CI-like proteins with only the NTD repressor.

### DNA-damaging agent triggers the induction of *Bacteroidales* prophage LoVE encoding a complete CI-like repressor

Given that Bt_806 carried six prophages, with only one encoding a complete CI-like repressor, we selected this isolate to delve into the transcriptional dynamics of prophage induction within a gut poly-lysogen. Based on our hypothesis that prophage LoVE, the only prophage within Bt_806 encoding a complete CI-like repressor, would respond to DNA damage, we tested whether the DNA-damaging agent, mitomycin C (MMC), triggered the induction of this phage. We first evaluated the effect of MMC on bacterial growth by comparing the optical density (OD_600_) of Bt_806 grown in control media (YCFA) and media supplemented with 0.3 µg/ml of MMC. Compared to control, Bt_806 exhibited impaired growth, with a classical sigmoid pattern that was indicative of prophage induction, roughly 4.5 h after the addition of MMC (**Fig. 2A**).

**Figure 2.**
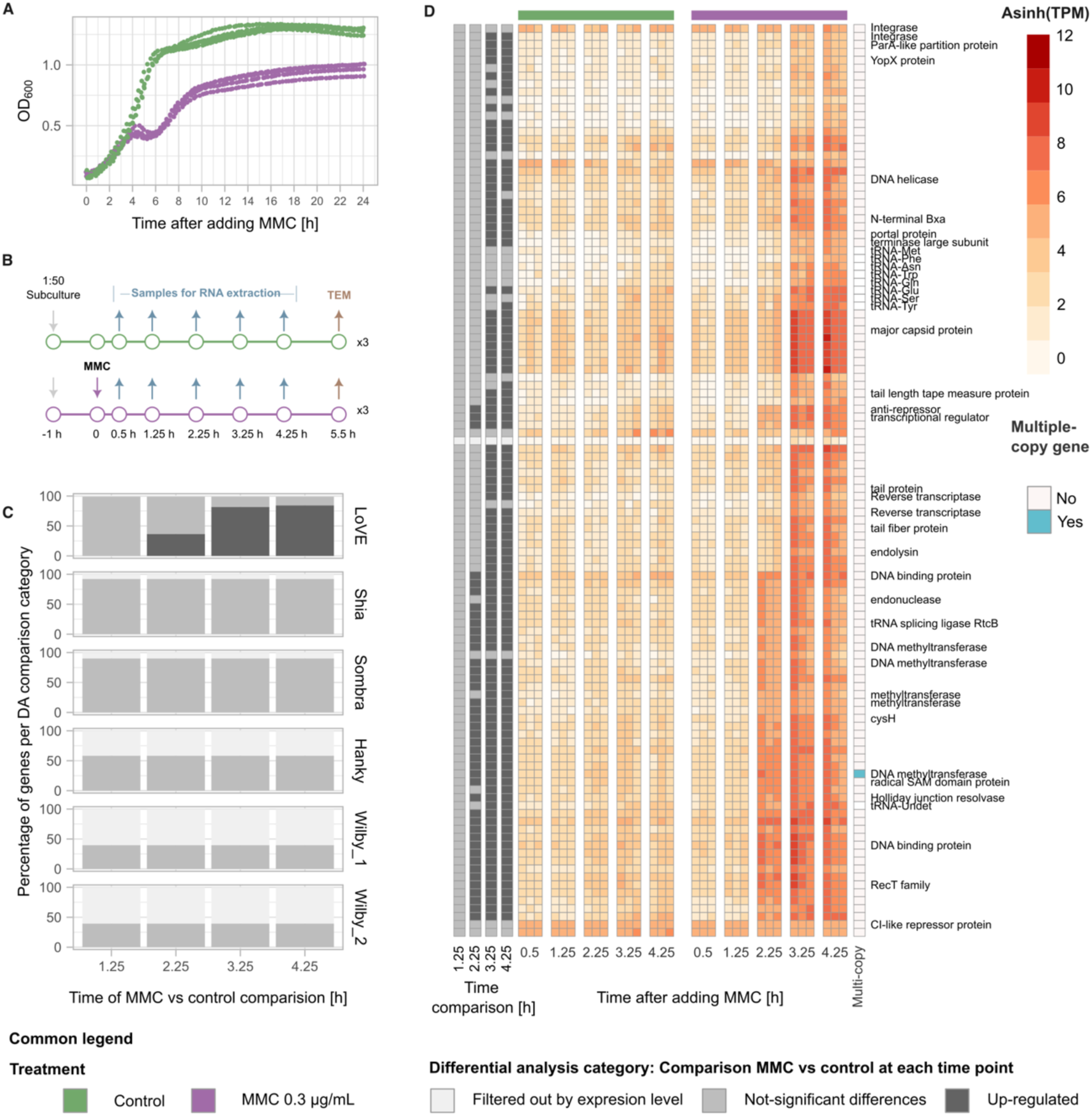
LoVE prophage was induced upon DNA damage. **A)** Growth curve of isolate Bt_806 grown in control media (green) and = media with 0.3 µg/mL MMC (purple). **B)** Experimental design to evaluate the transcriptional activation of prophage LoVE upon MMC induction. **C)** Categorization of prophage-encoded genes based on the expression level. Genes with low read count were filtered out of the differential analysis (light grey). The percentage of expressed genes with non-significant differences across treatments are shown in grey, and significantly up-regulated genes in MMC compared to control samples are shown in dark grey. **D)** Inverse hyperbolic sine of transcripts per million (TPM) of prophage LoVE genes in control media (green) and media with 0.3 µg/mL MMC (purple) are shown in the central heatmaps. Data from each biological replicate (n=3) is presented in separate columns at each time point. Columns in grey scale at the left of the heatmaps indicate whether a gene was differentially expressed or filtered out of the analysis at a specific time (same scale than panel C). The column at the right of the heatmaps highlights genes that have multiple copies across the Bt_806 genome. Functional annotation is presented at the right of this last column; hypothetical/unknown genes were left blank.

To validate prophage induction within Bt_806, we assessed the transcriptional response of Bt_806 cultures at 0.5, 1.25, 2.25, 3.25, and 4.25 h after adding 0.3 µg/mL MMC compared with a non-treated control. We extracted and sequenced RNA from each time point with three paired biological replicates per treatment (**Fig. 2B**). An additional sample was collected at 5.5 h following the addition of MMC for transmission electron microscopy (TEM) imaging. In addition to filtering out low-quality and human-contaminated reads, we excluded genes with low read counts for further analyses (filtered out by expression level category in **Fig. 2C**). More than 40% of Wilby 1, Wilby 2, and Hanky genes (60%, 61%, and 42%, respectively) were excluded from the analyses due to low read count. This observation, coupled with their transcriptional response (Supplementary Fig. 3), suggests that these three prophages were not induced during the experiment.

To evaluate the differential expression of control versus MMC-treated samples, we performed a paired differential analysis within each time point using the 0.5 h time point as the baseline. Among the six Bt_806 prophages, only LoVE presented differentially expressed genes between the treatments (**Fig. 2C**). Compared to the control, 42, 94, and 97 differentially expressed phage LoVE genes were up regulated 2.25, 3.25, and 4.25 h after the addition of MMC, respectively (**Fig. 2D**). Thus, upon DNA damage (i.e. after addition of MMC) the complete CI-encoding prophage LoVE was transcriptionally induced and followed a temporal dynamic in which genes were organised and expressed in clusters of early and middle-late genes, as has been observed in other phages^37,38^.

Several of the up-regulated prophage LoVE genes provided insights into prophage life cycles and their potential impact on their bacterial host. As such, the early expression of *cysH*, a potential auxiliary metabolic gene associated with cysteine biosynthesis^39^, suggests a role in nutrient acquisition as part of host takeover. Additionally, the early expression of genes encoding an anti-repressor protein and a transcriptional regulator, which, given their proximity to lytic genes, such as the tail length tape measure protein and the major capsid, we hypothesise were involved in the commitment to the lytic cycle by activating the expression of the middle-late genes. We also observed an early up-regulated array of genes encoding four methyltransferases, a tRNA splicing ligase RtcB, and an endonuclease. This array has been found in other members of the candidate family “Quimbyviridae”, to which LoVE phage belongs, and was proposed to function in counter-defence mechanisms relevant in phage-host infection^33^. The up-regulation of middle-late LoVE genes included an endolysin, tRNAs, and virion assembly proteins, such as terminase, major capsid protein, tape measure protein, and portal protein. Thus, the early expressed LoVE genes were broadly associated with the suppression of lysogeny, transcriptional activation of middle/late genes, DNA and RNA metabolism, host takeover, and counter-defence mechanisms, while middle-late genes pivot towards LoVE virion propagation, assembly, and eventual host cell lysis.

In terms of differentially expressed bacterial genes, we found three and six bacterial genes that were up-regulated 3.25 and 4.25 h after adding MMC, respectively. Three of these differentially expressed bacterial genes were located upstream and in the opposite direction to a complete CI-like repressor encoded within the Bt_806 genome, which was not associated with an intact prophage and is suggestive of a broader role of these transcriptional repressor domains in *Bacteroidales* genomes^40^ (Supplementary Fig. 4).

### Bt_806 prophages are transcriptionally active during lysogeny and contribute to prophage maintenance

To further explore the biology of Bt_806 temperate phages during lysogeny, we delved into prophage-encoded genes that were constitutively expressed (TPM >=10 across all samples and time points) without significant differences between treatments. We found that all Bt_806 prophages constitutively expressed clusters of genes across all time points and samples (**Fig. 3A)**, which included repressors, toxin-antitoxins (TA) systems, and genes of unknown function (Supplementary Fig. 5). Interestingly, the CI-like repressors within LoVE, Sombra, Shia, and Hanky were co-expressed alongside one or more adjacent hypothetical genes (**Fig. 3B**). In Shia and Sombra, these CI-like-adjacent hypothetical genes encoded proteins with either signal peptides, outer membrane, and/or transmembrane domains, which suggests novel induction mechanisms that may be associated with inter-cell phage-to-phage communication, similar to the described arbitrium systems that senses viral abundance as proxy for host availability^41^.

**Figure 3.**
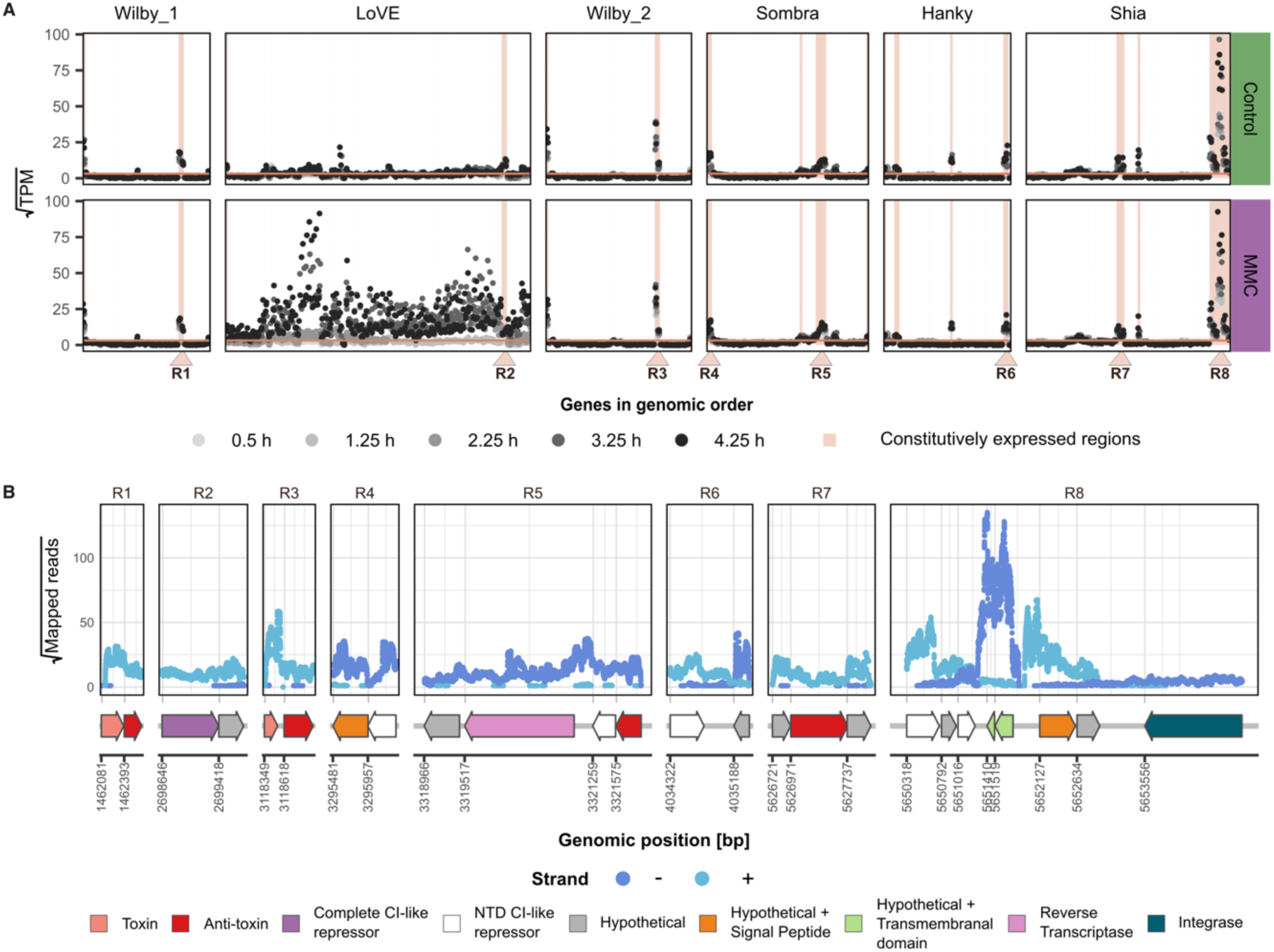
*Bacteroidales* prophages are transcriptionally active during lysogeny. **A)** Transcripts per million (TPM) of genes encoded by Bt_806 prophages in control media (green) and media with 0.3 µg/mL MMC (purple) across time points (points gray scale, n= 3 per time point per treatment) are shown in their genomic order. Regions of genes with an expression value higher than 10 (horizontal orange line) across all time points are shaded, and eight regions of interest (R1 to R8) were highlighted. **B)** Read depth across the eight regions of interest is presented. Given there were no differences across time points and samples, each point represents a sample from either the control or MMC-treated treatments at only 4.25 h (n=6 per base pair, bp). Only data with a read depth higher than zero is presented.

Putative toxin-antitoxin (TA) systems were constitutively expressed in Sombra, Shia, and the two Wilby prophages. In contrast to classical TA II systems, which are two-gene operons that encode a toxin and its cognate antitoxin^42^, Sombra and Shia TA genes were co-expressed with additional adjacent genes (**Fig. 3B** and Supplementary Fig. 5). Sombra, which encoded the HigB toxin, also expressed a reverse transcriptase and a hypothetical gene with unknown function, while Shia only expressed its antitoxin (an ADP-ribosylglycohydrolase) and two adjacent hypothetical genes. Variation of the HigBA system where more than one gene is co-expressed has been shown to provide broad-spectrum antiphage defence^43^. Thus, we hypothesised that Sombra and Shia TA systems, which were co-expressed with additional adjacent genes of unknown function, may have roles beyond maintenance of these prophages within this host.

The co-infection of Bt_806 with two near-identical Wilby prophages, Wilby 1 (44.3 K bp) and Wilby 2 (45.6 K bp), is compelling (Supplementary Fig. 6A). Co-infection with the same prophage has been previously observed^44,45^ and was associated with novel excisionases and mutations in the repressor. Here, the two Wilby prophages were integrated into different genomic regions and encoded distinct integrases and NTD CI-like repressors. While their integrases did not share amino acid identity, their NTD CI-like repressors shared 61% amino acid identity. In addition, they each encoded a different, putative two-gene TA II system that was constitutively expressed (**Fig. 3B**). Two-gene TA II systems were first reported on plasmids as “addiction modules”, which maintain plasmid stability by eliminating progeny cells that failed to inherit the TA-encoding plasmid^46^. We hypothesized that these distinct TA II systems may facilitate long-term stability and co-existence of the two near-identical Wilby prophages. To validate this, we performed a 15-day experiment consisting of daily passages in triplicate of isolate Bt_806, followed by colony PCR of both Wilby prophages at days five, ten, and fifteen of the experiment (Supplementary Fig. 6B-C). Both Wilby prophages were found in all isolates tested (*n*=120), indicating their stability within Bt_806. To validate the function of the Wilby TA II systems in prophage maintenance, we separately cloned and expressed the toxin and the complete TA system of each Wilby phage into *E. coli* DH5α. We confirmed that each toxin had a negative effect on *E. coli* growth, with the HicA toxin having a stronger effect than the DhiT toxin (Supplementary Fig. 6D). When the complete TA systems were expressed, the DhiTA system^47^ (Wilby 1) had negligible impact on growth, while the HicAB system (Wilby 2) did not entirely rescue host growth compared to control (Supplementary Fig. 6D), suggesting that other *Bacteroidales* proteins may be necessary to neutralize the effect of the HicA toxin in the native system. Together, these results demonstrated that Bt_806 prophages are transcriptionally active during lysogeny, express clusters of genes that are involved in repressing the lytic cycle and maintenance of prophages in their bacterial host progeny.

### Prophages LoVE and Sombra were spontaneously induced

We showed that prophage LoVE, which was the only prophage in isolate Bt_806 to encode a complete CI-repressor, was transcriptionally induced following the addition of MMC. To corroborate transcriptional induction with the release of LoVE virions, we collected both control and MMC-treated samples 5.5 h after addition of MMC, which were visualised with TEM. Surprisingly, we detected two structurally distinct viral-like particles (VLPs) in MMC-treated and control samples. Within the MMC-treated samples, we observed a larger-sized VLP with a tail length of 315 ± 11 nm (*n*=15), which we speculate is phage LoVE. Within the control samples, we visualised a smaller VLP with a tail length of 247.1 ± 9.1 nm (*n*=15) (**Fig. 4A**), suggesting spontaneous induction of a separate prophage.

**Figure 4.**
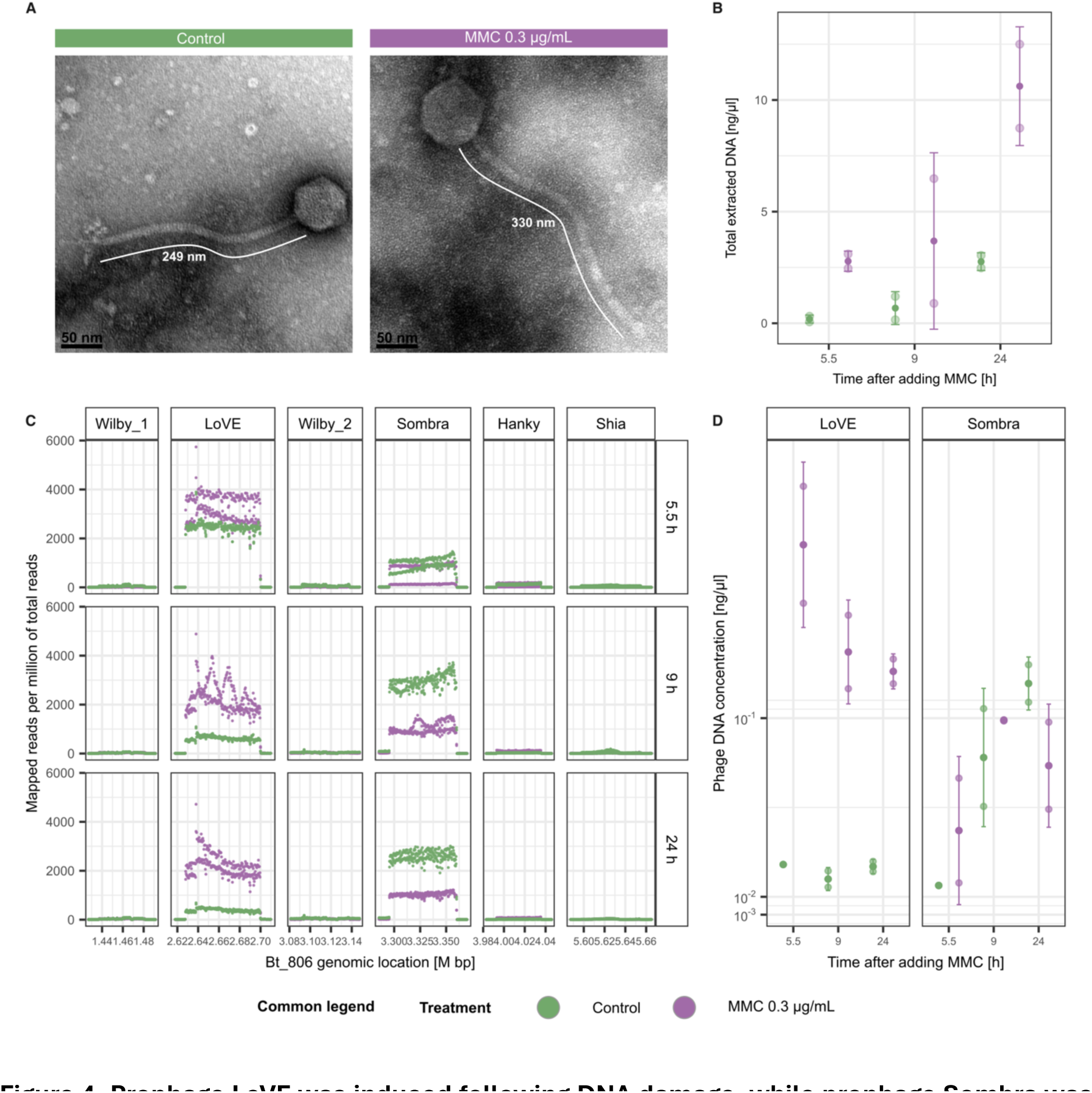
Prophage LoVE was induced following DNA damage, while prophage Sombra was spontaneously induced. **A)** Visualisation of smaller-sized virus-like particles (VLPs) in control samples (tail length 247.1 ± 9.1 nm, n=15, green bar) and larger-sized VLPs in MMC-treated samples (tail length 315.7 ± 11.6 nm, n=15, purple bar). **B)** To validate the source of these virions, DNA sequencing and absolute quantification through qPCR from enriched viral-like particles (VLPs) of control (green) and MMC-treated (purple) samples was performed. Samples from two replicates were collected at 5.5, 9, and 24 h after adding MMC. The concentration of total DNA extracted from these VLPs is presented. **C)** Profiles of VLP-sequenced reads mapped to Bt_806 prophage regions are shown. **D)** Absolute qPCR quantification of prophages LoVE and Sombra are shown. The dark dots represent the mean of the biological replicates, the whiskers represent the 95% CI, and the translucent dots represent the mean of three technical replicates performed during the qPCRs.

To verify spontaneous induction of Bt_806 prophages and confirm MMC-induced LoVE phage propagation, we enriched and sequenced VLPs at 5.5, 9, and 24 h across both control and MMC-treated samples. Compared to control samples, the concentration of DNA extracted from VLPs in MMC-treated samples was higher across all time points and replicates, indicating a higher concentration of induced virions following MMC-treatment (**Fig. 4B**). To verify the source of these virions, we sequenced and mapped the obtained reads to Bt_806 genome, observing an enrichment of reads mapping to prophages LoVE and Sombra across all time points from both control and MMC-treated samples (**Fig. 4C**). We then quantified the abundance of Bt_806 LoVE and Sombra virions using quantitative PCR (qPCR) from the same virion-sequenced samples. Aligned with our prior observations, we detected an average of 7.1 times more phage LoVE DNA at the 5.5 h time point in MMC than control samples (**Fig. 4D**), with similar increases of 7.0 and 4.9 in LoVE DNA concentration compared to the control at 9 and 24 h, respectively. Looking at prophage Sombra, we detected a 1.5 times lower average concentration of virion DNA in MMC treated compared to control samples after 24h (**Fig. 4D**). Although there was an increase of Sombra virions in MMC-treated samples compared to control after 5.5 h (2.8 times higher) and 9 h (1.2 times higher) (**Fig. 4D**), the increase was not as high as the increase with LoVE virions, suggesting that Sombra phage was spontaneously induced regardless of the addition of MMC. Together, these results suggest that MMC-mediated induction led to the release of prophage LoVE virions, but spontaneous induction of both LoVE and Sombra virions occurred at lower levels in the complex YCFA media.

### Diverse complete-CI *Bacteroidales* prophages are induced in response to DNA damage

Our initial screening identified 197 complete-CI *Bacteroidales* prophages, suggesting that these prophages may be inducible in response to DNA damage. Consistent with this, MMC exclusively triggered induction of prophage LoVE in the poly-lysogen Bt_806, supporting the use of these genes as markers to predict DNA-damage-inducible prophages within gut lysogens. To test this, we selected an additional eleven *Bacteroidales* poly-lysogens, each carrying a unique prophage composition, yet with only one prophage in each isolate encoding a complete CI-like repressor (**Fig. 5A**). Employing these eleven poly-lysogens, we tested the induction of prophages belonging to five different VCs (LoVE, Pomma, VC_217, VC_156, and VC_126). Given the high prevalence of prophages LoVE and Pomma, their induction consistency was further assessed across multiple hosts. As before, we first assessed the growth curve of these eleven isolates, observing reductions in OD and a classical sigmoid pattern consistent with prophage induction following the addition of the DNA-damaging agent MMC, although the timing and MMC concentration varied among strains (Supplementary Fig. 7). To quantify induction, cultures were grown under control and MMC-treated conditions, viral-like particles were isolated, and induced complete-CI prophages were quantified by qPCR, with sampling times and MMC concentrations optimized for each isolate; Bt_806 was included as a positive control. Across all isolates, MMC treatment resulted in higher levels of DNA from complete-CI prophages in virion fractions compared to controls, with nine showing statistically significant differences (adjusted p-value < 0.05; **Fig. 5B**). Together, these results demonstrate that induction of *Bacteroidales* prophages encoding complete CI-like repressors was consistently triggered by DNA damage across diverse viral clusters and host genetic backgrounds. In summary, this demonstrates the capacity to informatically identify prophage repressors within *Bacteroidales* hosts, which, alongside a mechanistic understanding of their induction, enables the use of phage repressor genes as a reliable marker of prophage induction in the gut microbiome.

**Figure 5.**
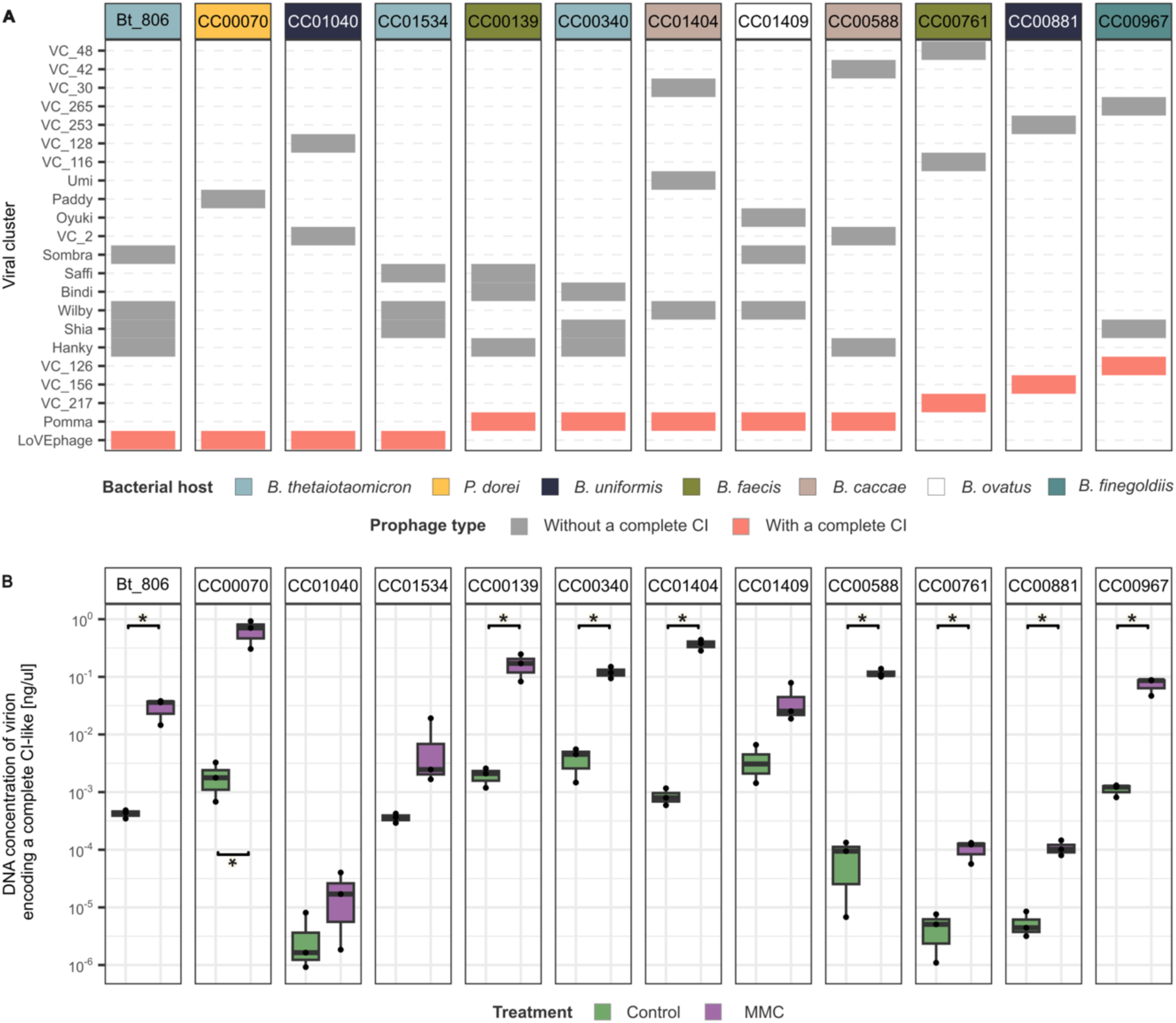
*Bacteroidales* complete-CI prophages are induced upon the addition of a DNA-damaging agent. **A)** Eleven poly-lysogens from two *Bacteroidales* genera (represented in the colour of each column), which harboured a unique prophage composition, each comprising only one complete-CI prophage (salmon tiles). Bt_806 was included as a positive control. **B)** Isolates were induced, and the absolute DNA concentration of virions encoding a complete CI-like repressor (qPCR) was quantified in both control (green) and MMC-treated (purple) samples. Three biological replicates were employed per isolate. *: FDR <0.05, paired t-test. Black points represent the mean of three technical qPCRs performed for each biological replicate.

## Discussion

Bacterial species of the *Bacteroidales* order largely colonise and impact the human gut ecosystem^48^. Prophages are prevalent within gut *Bacteroidales* species^8^; in our collection, ∼91% of the *Bacteroidales* isolates carried at least one prophage, and ∼ 72% carried two or more. The spontaneous induction of these prophages is predicted to occur at low rates within the gut of healthy human adults^7,49^ and increase at specific life-stages or disease states. Thus, the high prevalence of *Bacteroidales* prophages in the human gut and their life stage-associated induction highlights the need to define the genetics and mechanisms underlying the lysis-lysogeny decision. Based on the current understanding of the lysis-lysogeny switch in phage Lambda, we explored a collection of 335 *Bacteroidales* isolates that harboured 902 prophages to identify complete CI-like repressor proteins as genetic regulators of induction. Despite extensive evolutionary divergence in these CI-like repressors, we demonstrate their identification in 197 *Bacteroidales* prophages through an iterative profile Hidden Markov Model. We then validate that *Bacteroidales* prophages encoding such repressors were predictably induced in response to DNA damage. This response was consistently observed across prophages encoding complete CI-like repressors from five different viral clusters, and in the case of the promiscuous *Bacteroidales* prophages LoVE and Pomma – which both encode complete CI-like repressors – the response was consistent across distinct bacterial-host genetic backgrounds. Together, these findings demonstrate that core regulatory gene components of lysogeny extend to dominant gut-associated bacteria and suggest that we can identify and selectively control prophage induction by understanding the biology and genetics of these systems.

A bacterial lysogen harbours an integrated prophage within its chromosome, where the prophage encodes its own repressor module to control the lytic and lysogenic cycles^21^. However, most bacterial hosts, especially within the gut, are poly-lysogens that carry multiple prophages. Co-habiting prophages often encode diverse repressors and regulatory gene circuits, which can enable prophage crosstalk^25^. Thus, poly-lysogenic hosts are fascinating systems to understand how their integrated prophages coordinate induction, repress co-inhabiting phage functions, and ultimately compete for the limited host cellular resources available to each prophage to complete their replicative cycles. We focused on the poly-lysogenic *Bacteroides thetaiotaomicron* isolate Bt_806, which carries six prophages, only one of which encoded a complete CI-like repressor. Using this model, we characterized the temporal transcriptional dynamics underlying the lysogeny-to-lytic switch of prophage LoVE, which was induced upon DNA damage, leading to the transcriptional activation of early, middle-late genes associated with the lytic cycle.

Consistent with the emerging view that prophages are not merely inactive elements awaiting induction^5^, we show that resident *Bacteroidales* prophages are transcriptionally active during stable lysogeny, expressing multiple distinct gene clusters with putative functional impacts on their bacterial hosts. As expected, constitutively expressed genes included the CI-like repressors, supporting their role in regulating the lytic switch. In addition to regulatory genes, we identified constitutively expressed gene clusters encoding toxin–antitoxin (TA) systems. We provide evidence of two toxin genes, *hicA*and *dhiT*, expressed by the co-inhabiting and highly similar Wilby phages that were likely associated with prophage addiction and maintenance.

The evolution of complex regulatory circuits, such as the lysis-lysogeny genetic switch, has been proposed to follow a two-stage model where an early gene circuit evolved from a primitive or basic form that was subsequently refined through the addition of new features^21^. Thus, identifying the most basic form of prophage-induction circuits could provide a principled framework for predicting how and when gut prophages respond to environmental cues across diverse hosts. Importantly, the predictive power of CI-like repressors does not rely on close nucleotide homology, but rather on the retention of key functional domains, underscoring the modular nature of prophage regulatory circuits. Consistent with this, our results link the presence of complete CI-like repressors to DNA damage-driven induction of *Bacteroidales* prophages. Further, we identified hundreds of *Bacteroidales* prophages that lack the CI-like CTD self-inactivator, which is required for self-cleavage in response to DNA damage, underscoring the potential diversity of novel, DNA-damage-independent prophage induction mechanisms that remain to be characterised.

Alternative prophage induction strategies have recently been described in other systems, including *Bacillus* phages that employ small “arbitrium” peptides to communicate and coordinate lysis-lysogeny decisions^50,51^. Analogously, prophage Shia in our isolate Bt_806 constitutively expressed a gene encoding a signal peptide that was adjacent to its CI-like repressor^52^, raising the possibility that *Bacteroidales* prophages also engage in peptide-mediated phage communication. Together, these lysogeny-associated gene clusters provide a foundation for future functional characterization of hypothetical genes that may mediate previously unrecognized phage–phage or phage–host interactions in the gut microbiome. A deeper understanding of prophage induction and their associated regulatory genetic architectures may ultimately enable the selective induction of gut prophages, offering new avenues to control prophage induction and modulate gut microbiome dynamics.

While we were able to link the presence of a complete CI-like repressor with DNA-damage-driven induction of *Bacteroidales* prophages, this approach has several limitations. First, prophage induction may be intrinsically dependent on host physiology. As such, experimental parameters, including the selected growth media, *in vitro* conditions, the concentration of MMC, the timing of its addition, and the sampling window relative to lysis may impact our ability to detect prophage induction. Consequently, the absence of induction under a given experimental condition does not necessitate the inability of a prophage to respond to a given cue. Second, co-habiting prophages within the same poly-lysogen may encode competitive regulatory circuits aimed to inhibit the induction of other prophages, or they may encode circuits that sense and respond faster, ensuring their propagation over prophages encoding complete CI-like repressors. Third, the bacterial host may encode defence systems to stall the induction of the prophage. Together, these considerations highlight that complete CI-like repressors provide a useful but context-dependent framework for predicting prophage induction and motivate future efforts to incorporate alternative regulatory modules and physiologically relevant cues to better resolve prophage dynamics in complex microbial communities. Future work will be required to identify alternative induction cues, elucidate the regulatory mechanisms of prophages lacking complete CI-like repressors, compare prophage induction *in situ*, and determine the functional roles of constitutively expressed lysogeny-associated genes of unknown function. Nevertheless, by combining predictive genomics with experimental validation, our work provides a conceptual and methodological framework for dissecting prophage regulation in complex microbiomes and emphasizes that prophages function as active genomic elements rather than dormant passengers.

## Data availability

Reads obtained from the RNA-seq experiment were deposited in the European Nucleotide Archive (ENA) at EMBL-EBI under project number PRJEB110793. Bacterial isolates are available through the Australian Microbiome Culture Collection (AusMiCC). Bacterial genomes from AusMiCC are also stored in ENA, and their accession number are provided in Supplementary Table 2. *Bacteroidales* prophage predicted genomes and bioinformatic scripts are available at https://github.com/luayou/prophage_induction.git.

## Acknowledgements

This research was supported by the Australian Research Council (ARC) Discovery Project Grant (DP260102966); J.J.B was supported by a National Health and Medical Research Council (NHMRC) Investigator Grant Leadership Level (IG2026130). Monash eResearch capabilities supported the infrastructure, including high-performance computing, M3, and vault tape-based storage, to analyse and store the data. We thank the Knott lab for providing us with the pTET plasmid, specifically to Giovanni Leandri, who guided us in the design of the constructs. Additionally, thanks to Alexandra McAllan for helping us to upload the bacterial genomes to ENA.

## Author contributions

L.A-F, S.C.F, and J.J.B conceived and designed the study. L.A-F performed all the experiments within this manuscript, including *in vitro* inductions, molecular work, RNA extractions, and informatics analyses. S.D contributed ideas and expertise. E.L.R assisted with the accession and information referred to the AusMiCC isolates. R.B.Y obtained Bt_806 DNA to long-read sequencing. J.A.G conducted all library preparation and sequencing. D.K. conducted TEM grid preparation and TEM imaging. S.C.F, and J.J.B funded the research. L.A-F and J.J.B drafted the original manuscript. All authors contributed to the review and editing of this manuscript.

## Methods

### The gut bacterial genomes

This study employed 989 bacterial genomes obtained from the Australian Microbiome Culture Collection (AusMiCC). The collection comprised genomes from 127 *Actinomycetota*, 353 *Bacillota*, 335 *Bacteroidota*, 11 *Fusobacteriota*, and 163 *Pseudomonadota* gut-derived isolates (Supplementary Table 2). Confirmation of AusMiCC taxonomic assignment of *Bacteroidales* genomes was conducted using GTDB-Tk v. 2.4.0^35^ based on release 226 of their database^53^ (Supplementary Table 1).

### Bacterial culture conditions

Twelve *Bacteroidales* isolates obtained from the AusMiCC (CC00806, CC00070, CC01040, CC01534, CC00139, CC00340, CC01404, CC01409, CC00588, CC00761, CC00881, and CC00967) were employed to conduct wet-lab experiments in this study. Cultures of these isolates were grown in yeast-extract-casitone-fatty acids (YCFA) media^54^ at 37 °C under anaerobic conditions containing 10% carbon dioxide, 10% hydrogen, and 80% nitrogen (Whitley A95 or Whitley A85 anaerobic workstation). Frozen stocks of each isolate were maintained in glycerol suspension (25% v/v) at -80 ℃. For each experiment, isolates were streaked out onto YCFA agar plates and incubated for 24 h.

### Prediction of prophage regions and functional annotation

To predict prophages, we followed a modified protocol of Guo et al. ^55^. As such, we first predicted prophages using VirSorter2 v2.2.4^28^ (--keep-original-seq --min-length 5000 --min-score 0.5 --include-groups dsDNAphage, ssDNA). We then employed CheckV v0.7.0 based on checkv-db-v1.5^29^ to calculate the completeness of each predicted prophage and detect potential bacterial-host contamination at the flanked regions of the original predicted prophage. The flanked regions of prophages with potential bacterial-host contamination were trimmed, and their completeness was re-evaluated using CheckV ^29^. Trimmed prophages were selected over the original ones when their completeness was higher than or equal to the completeness of the original one. We found some trimmed prophage regions lacked integrase or transposase genes located at the end of the original prophage-region; in this event, up to 15 K bp at each end of the trimmed prophages (without exceeding the original predicted prophage region) were added to predicted genomes using bedtools v.2.30.0^56^. All predicted regions were then grouped at 95% average nucleotide identity (ANI) over 30% of the alignment fraction using scripts from the checkV repository. We then filtered out clusters of prophages that belong to ANI-clusters where none of the prophages have at least 50% genome completeness. The selected prophages were clustered into Viral Clusters (VCs) at 95% average nucleotide identity (ANI) over 85% of the alignment fraction, adding prophages from Dahlman et al.^8^ study to annotated pre-characterised prophages. Prophages belonging to the same VC within the same host were considered as a single prophage.

The selected prophage genomes were then annotated using DRAM-v v1.5.0^30^ coupled with VirSorter2 (--seqname-suffix-off --viral-gene-enrich-off --prep-for-dramv --min-length 5000 --include-groups dsDNAphage,ssDNA all). Based on the functional annotation, we performed manual curation exclusively of *Bacteroidales* prophages. For this, we included *Bacteroidales* prophage genomes from Dahlman et al.^8^ and Nayfach et al.^57^ studies and grouped them at 95% average nucleotide identity (ANI) over 30% of the alignment fraction with the *Bacteroidales* predicted prophage regions. Prophage genomes from Dahlman et al.^8^ were employed as references to trim other prophages from the same ANI at 30% coverage cluster. The trimmed prophages were then clustered with Dahlman et al.^8^ prophage genomes at 95% average nucleotide identity (ANI) over 85% of the alignment fraction, and a second round of manual trimming was performed. Completeness of the twice-trimmed prophages was recalculated using CheckV. Only prophages with at least a 40% complete genome belonging to VCs where at least one prophage has at least a 50% complete genome were kept for further analyses. As a result, we obtained 902 manually trimmed *Bacteroidales* prophages (Supplementary Table 3) and 1,271 prophage-regions from the remaining four bacterial phyla (Supplementary Table 4).

### CI-like repressor iterative search

We sought Lambda CI-like repressor proteins in a collection of 902 *Bacteroidales*prophages and 1,271 additional prophage regions predicted from *Actinomycetota*, *Bacillota*, *Pseudomonadota*, and *Fusobacteriota* isolates. For this, we employed the Lambda CI repressor (UniProtKB entry P03034) as a seed for a profile Hidden Markov Model search (profile-HMM). The search comprised seven iterations and was conducted using the HMMER package v.3.4 ^58^ (jackhmmer -N 7 --incE 1e-10 --incdomE 1e-10). Only hits with a bit score higher than 30 that cover at least 70% of the protein were defined as CI-like repressor proteins from the domain hits table. Hits covering from position three to position 193 of the profile-HMM were considered as complete CI-like proteins; hits covering from position three to any location between positions 92 and132 were considered as CI-like repressors with only the DNA-binding N-terminal domain (NTD CI-like repressors). Proteins’ IDs considered as complete or NTD CI-like repressor proteins can be found in Supplementary Table 3 and Supplementary Table 4.

### Bacterial phylogenetic tree

A phylogenetic tree was built de novo from protein sequence alignments of 120 core bacterial genes generated by GTDB-Tk v. 2.4.0^35^ based on release 226 of their database^53^. The tree was then reconstructed using IQ-TREE v.2.0.rc1, where LG+F+I+G4 was selected as the best substitution model. The tree was visualised and annotated in the frame of the iTOL online server^59^.

### Phylogenetic tree reconstruction of complete CI-like Lambda repressor proteins

A total of 417 complete CI-like repressor proteins encoded by gut prophages were dereplicated into 133 clusters at 100% amino acid identity over 100% coverage of the short one using CD-HIT v.4.8.1 (-c 1 -n 5 -g 1 -d 0 -G 0 -aS 1)^60^. Representative amino acid sequences from these clusters were aligned to the Lambda CI-like repressor protein using MAFFT v. 7.526^61^ (L-INS-I algorithm with default settings). Alignments were cleaned using the CIAlign v.1.1.4^62^ command-line tool (adding these two settings: --remove_divergent_minperc 0.1 --sequence_logo_type both to the default parameters). The final alignment was employed to reconstruct the phylogenetic tree using IQ-TREE v.2.0-rc1^36^ with the following settings: -m MFP -b 1000 -nt AUTO. The VT+F+R6 model was chosen to reconstruct the tree according to the Bayesian Information Criterion.

### Long-read sequencing

Single colonies of *Bacteroides thetaiotaomicron* CC00806 (Bt_806) were grown overnight in 40 mL of pre-reduced YCFA media, pelleted by centrifugation at 4,000 g for 10 min, and washed four times in 1 mL of PBS. DNA was extracted using the Monarch HMW DNA extraction kit (New England Biolabs) following the Gram-positive Bacteria protocol, with modifications as follows. Cells were lysed in 300 μL of STET buffer (8% sucrose, 5% Triton X-100, 50 mM EDTA, 50 mM Tris, pH 8) containing 10 mg/ml lysozyme, 300 μL of HMW gDNA Tissue Lysis Buffer, and 20 μL of Proteinase K, at 56 ℃ for 10 min. Lysates were treated with 10 μL of RNase A at 56 ℃ for 5 min, followed by 300 μL of Protein Separation Solution. Samples were mixed by inversion for 2 min, then centrifuged at 4 ℃ for 20 min at 16,000 g. Supernatants were collected, and 550 μL of isopropanol was added to 800 μL of supernatant. Samples were inverted for 5 min, or until DNA was precipitated, and DNA was pelleted by centrifugation for 10 min at 4 ℃ and 12,000 g. The resulting pellet was washed twice with 500 μL of gDNA wash buffer and resuspended in nuclease-free water. Library preparation and Oxford Nanopore MinION sequencing were performed using the Oxford Nanopore ligation sequencing kit with native barcoding expansion kit. The obtained Oxford Nanopore Technologies long reads and the available AusMiCC Illumina short reads were used to conduct a hybrid assembly using Dragonflye v1.1.1^63^. The assembled genome was annotated using DRAM in single mode.

### Stability of Wilby 1 and Wilby 2 prophages through bacterial generations

Three single colonies of Bt_806 were separately grown in pre-reduced YCFA media for 24 h. The cultures were diluted 1:50 in 3 mL of pre-reduced YCFA and incubated in anaerobic conditions at 37 ℃ for 24 h. Daily 1:50 passages were conducted into fresh pre-reduced YCFA media during five days. After five days, 60 μL of each culture were diluted into 60 μL of 50% glycerol and stored at 80 ℃ until the experiment was resumed. In addition to this, serial dilutions of each culture were conducted and plated out onto YCFA plates; eight single colonies were screened to evaluate the persistence of Wilby 1 and Wilby 2 through PCR (refer to Supplementary Table 5 for primer sequences). The experiment was resumed by reconstituting the whole glycerol stock into 3 mL of pre-reduced YCFA. Daily passage, sampling, and storage of the samples were performed every five days as mentioned previously. After a total of 15 daily passages, we screened for Wilby 1 and Wilby 2 in 24 single colonies via PCR.

### Effect of mitomycin C on *Bacteroidales* growth curve

Single colonies were grown in pre-reduced YCFA media for 24 h. Cultures were then diluted 1:50 in 1.8 ml YCFA and grown for 1 hour before adding mitomycin C (MMC). After addition of MMC, an 180 μL aliquot was transferred to a 96-well plate. Growth was monitored by OD_600_ readings every 5 min for 40 h on a Cerillo plate reader incubated anaerobically at 37 ℃.

### RNA-seq experiment

Two single colonies of Bt_806 were separately grown in pre-reduced YCFA media for 24 h. The cultures were then diluted 1:50 in 200 mL of pre-reduced YCFA. After one hour of incubation, MMC (final concentration 0.3 μg/mL) was added to one of the cultures, while the other was used as a control. Samples for RNA extraction were taken from both treatments at 0.5, 1.25, 2.25, 3.25, and 4.25 h after addition of MMC. A final sample of each treatment was collected after 5.5 h of adding MMC and cleaned for transmission electron microscopy. This experiment was repeated two additional times on two separate days. Collected samples for RNA extraction were immediately centrifuged for 15 min at 4 ℃ and 4,000 g. The pelleted cells were resuspended in 1 mL TRIzol and lysed via bead-beating (Lysis matrix E) twice for 40 seconds, with an intermediate step of cooling on ice for 2 min. After this, the samples were centrifuged for 10 min at 4 ℃ and 12,000 g. The supernatant was stored at -80 ℃ until all the samples were processed. The thawed samples were then treated with 250 μL of chloroform for 3 min, inverting them occasionally, followed by centrifugation for 15 min at 4 ℃ and 10,000 g. RNA in the clear aqueous phase of each sample was purified using RNeasy® Mini Kit (Qiagen). The kit protocol was followed from step two, including the on-column DNase digestion. Library preparation was performed following the Illumina Stranded Total RNA Prep, Ligation with Ribo-Zero Plus Microbiome kit. For this, 200 ng of RNA was used as input and underwent 14-15 cycles of amplification. Final libraries were quantified by Qubit and a pool made based on concentration. The quality of the final library pool was measured by Qubit, Bioanalyzer, and qPCR. For sequencing, 1000 pM of the library pool was clustered on a P2 NextSeq2000 run.

### Differentiallly expressed and constitutively expressed genes

Sequencing reads obtained from control and MMC-treated samples across five time points were trimmed using Trimmomatic v.0.38 (SLIDINGWINDOW:4:20 CROP:57 HEADCROP:11 MINLEN:30)^64^. Trimmed reads were aligned to human DNA (GRCh38.p14) using Bowtie v.2.5.4^65^; reads that aligned to human DNA were filtered out of the analysis. Cleaned reads were mapped to the Bt_806 genome using Rsubread v.2.12.3^66^. Gene counts were obtained using featureCounts^67^ function from Rsubread. Each time point consisted of three biological replicates per treatment. A principal component analysis (PCA) was performed to visualise the overall patterns in the variation of the dataset. PCA was conducted using the plotMDS function in R. Low-expression genes were filtered out by using the filterByExpr function (min.count=10, large.n=3, min.prop=1) from edgeR v.4.4.2^68^. Counts of the remaining reads were normalised using the normLibSizes function from edgeR. Normalised counts were modelled using a quasi-likelihood (QL) negative binomial generalized log-linear model (glmQLFit function). Significantly differentially expressed (DE) genes were identified through a QL F-test; only genes with FDR < 0.05 and the absolute value of log_2_FC > 2 were considered as DE genes. Constitutively expressed genes were defined as genes with no significant difference across treatments and with a count in transcript per million (TPM) higher than ten. Raw gene counts were divided by the gene length in kilobases to obtain reads per kilobase (RPK). Then the total sum of all RPK values was divided by one million to obtain the scaling factor. TPM values per gene were obtained by dividing the RPK gene value by the scaling factor.

### Transmission electron microscopy (TEM) images

Lysates from induced bacteria were concentrated using an Amicon with a 1000 kDa filter following the Brum^69^ protocol prior to imaging. A 400-mesh copper TEM grid (SPI Supplies, USA) with carbon-stabilized ultrathin (invisible on the water surface)^70^ formvar support film was placed (face side down) on a 10 µl droplet of phage suspension for 30 s and then dried using filter paper. Subsequently, the grid was placed (face side down) on a 10 µl droplet of 1% (w/v) water solution of uranyl acetate for 30 s and dried using filter paper. Finally, the grid was examined with a TEM (JEM-1400 Plus; Jeol, Japan) at an accelerating voltage of 80 kV.

### Evaluating spontaneous induction of prophages LoVE and Sombra

Four single colonies of Bt_806 were grown separately in pre-reduced YCFA media for 24 h. The cultures were then diluted 1:50 in 200 mL of pre-reduced YCFA. After one hour of incubation, MMC (final concentration 0.3 μg/mL) was added to two of the cultures, while the other two were used as a control. 5 mL samples were taken at 5.5, 9, and 24 h after addition of MMC from both control and MMC-treated samples to enrich viral-like particles (VLPs). For this, collected samples were immediately centrifuged for 15 min at 4 ℃ and 4,000 g. The supernatants were treated with 10% chloroform for 15 min and centrifuged for 15 min at 4,000 g. The aqueous phase was then treated with 10 μg/mL DNase and 0.01 volume RNase for 1 hour at 37 °C. To precipitate VLPs from the samples, 7% PEG 8000 0.3 M NaCl solution was added, followed by incubation for 18 h at 4 °C and centrifugation for 45 min at 14,000 g. The obtained pellets were resuspended in 50 μl of TE buffer. Phenol-chloroform-isoamyl alcohol (25:24:1) DNA extraction from VLPs was conducted following the Dahlman et al.^8^ protocol. The total obtained DNA was quantified using Qubit. Nextera-XT libraries were then constructed from each DNA sample and sequenced on the NextSeq 2000 platform. After library preparation, quantification of absolute DNA from prophages Sombra and LoVE was conducted using quantitative PCR (qPCRs) (refer to Supplementary Table 5 for primer sequences). qPCRs were performed in technical triplicate using SYBR Green I Master kit containing 0.5 μM of each primer, 1.5 μL of DNA template, and 1X SYBR Green I Master mix, in a final reaction volume of 20 μL. Cycle parameters were as follows: initial denaturation at 95 °C for 10 mins; followed by 45 cycles of 95 °C for 10 sec, 62 °C for 20 sec, and 72 °C for 20 sec. To verify the specificity of the reaction, melt curves were performed after amplification. The standard curve was produced from 1 ng/μL to 1e-7 ng/μL using a 10 ng/μL gBlock containing Sombra or Love amplicon sequence (Supplementary Table 6). A linear model was built based on the standard curve of each primer and employed to calculate the absolute DNA of prophages LoVE and Sombra in the test samples. Given the low amount of DNA from one of the control replicates after 5.5 h of addition of MMC, this sample was not processed for quantifications of prophages Sombra and LoVE.

### Heterologous cloning and expression of TA systems in *Escherichia coli*

The native DNA locus of the toxins and the complete TA systems of each Wilby were amplified, adding BsbI enzyme-flanked regions from Bt_806 (refer to Supplementary Table 5 for primer sequences). Using Golden Gate assembly, the amplified regions were inserted under an anhydrotetracycline (aTc)-inducible promoter into a pTET vector carrying a chloramphenicol resistance marker. Cells of *E. coli* DH5α were then heat-shock transformed with each of the constructs. As a control, an empty plasmid was also transformed into *E. coli* DH5α cells. Transformants carrying each construct were maintained in glycerol suspension (25% v/v) at -80 °C. Transformant cells were streaked onto LB agar supplemented with 25 μg/mL chloramphenicol. Single colonies were grown at 37 °C in LB medium supplemented with 25 μg/mL chloramphenicol for 16 h. Cultures were diluted 1:50 into 25 mL of LB and grown until they reached an OD_600_ of 0.5. Four cultures were performed per transformant. After this, cells were collected and resuspended in fresh media with 100 nM aTc and 25 μg/mL chloramphenicol. An 180 μL aliquot of the resuspended cells per culture was transferred to a 96-well plate. Aerobic growth at 37 °C was monitored by OD_600_ readings every 5 mins for 24 h. Primers outside the inserts were designed (Supplementary Table 5) to verify the direction and sequence of the constructs using Sanger sequencing.

### Validating the induction of prophages encoding a complete CI-like repressor across five different viral clusters

Eleven additional poly-lysogens were employed to validate the induction of prophages encoding a complete CI-like repressor. Growth curves of these eleven isolates were obtained to determine the concentration of MMC and the sampling time for each isolate. The induction test was performed in triplicate in 1.8 mL of pre-reduced YCFA. As in previous experiments, this experiment started with growing single colonies in pre-reduced YCFA media for 24 h, followed by a 1:50 dilution into 1.8 mL of fresh media. One hour after the subculture, MMC at the defined concentration was added to the media. Samples without MMC were used as controls. After the defined sampling time, samples were collected for VLPs enrichment, DNA extraction, and qPCR quantification of each of the prophages encoding a complete CI-like repressor protein (refer to Supplementary Table 5 and Supplementary Table 6 for primer and gblocks sequences).

### Data analyses

All data analyses, including statistical analyses and visualization, were conducted using R v.4.4.2^71^. To organise and summarise the data, tidyverse v.2.0.0 and dplyr v.1.2.0 R packages were employed. edgeR v.4.4.2^68^ package was used to find differentially expressed genes (DEG); criteria to define DEG were mentioned before. Shapiro-Wilk’s test and Levene’s test were performed to check the normal distribution of the residuals and the homogeneity of variances of datasets obtained from qPCRs. A paired t-test was employed to compare phage DNA concentration of MMC-treated to control samples. False discovery rate (FDR) was used to adjust p-values from multiple comparisons; only comparisons with an FDR < 0.05 were deemed as significant. Locally weighted scatterplot smoothing (LOESS) regression was used to fit the trend of *Escherichia coli* DH5α growth over time. Raw plots were obtained using ggplot2 v.4.0.2, gggenes v.0.6.0, ggpubr v.0.6.3, or/and RColorBrewer V.1.1-3 R packages. Illustrations were made using Inkscape.

## Supplementary Figures

**Supplementary Figure 1.**
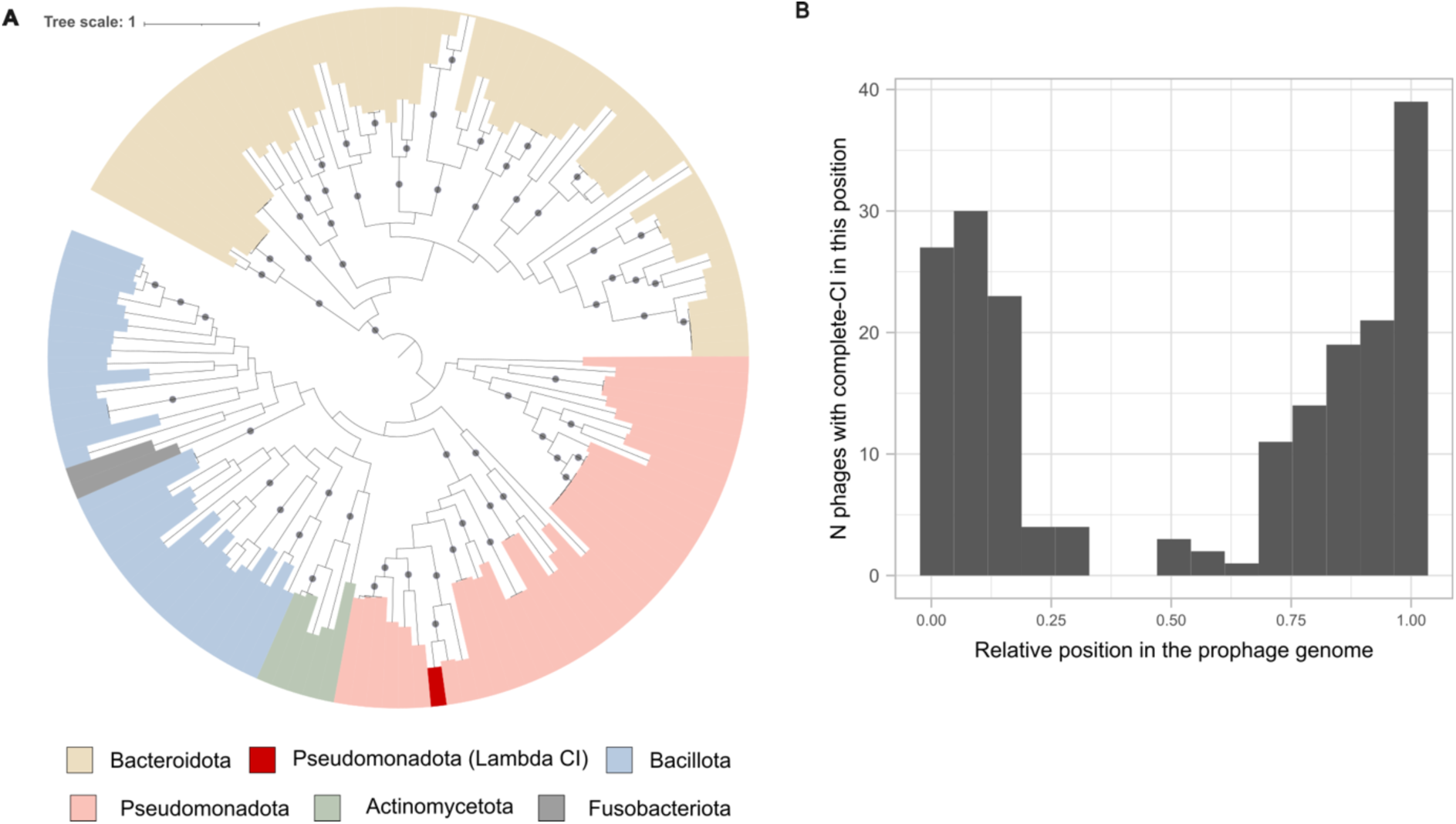
Prophage-encoded complete CI-like repressor phylogeny and relative position in prophage-genomes. **A)** Complete CI-like repressors were dereplicated at 100% amino acid identity over 100% of the length of the shorter sequence, and representative sequences were employed to reconstruct the tree. The internal black dots indicate conservative Bootstrap values higher than 80. While terminal clades are well supported, the topology of ancient splits is uncertain. **B)** Histogram of the number of prophages with a complete CI-like protein in each relative bin position to the prophage genome (n= 413 total prophages with a complete CI-like protein).

**Supplementary Figure 2.**
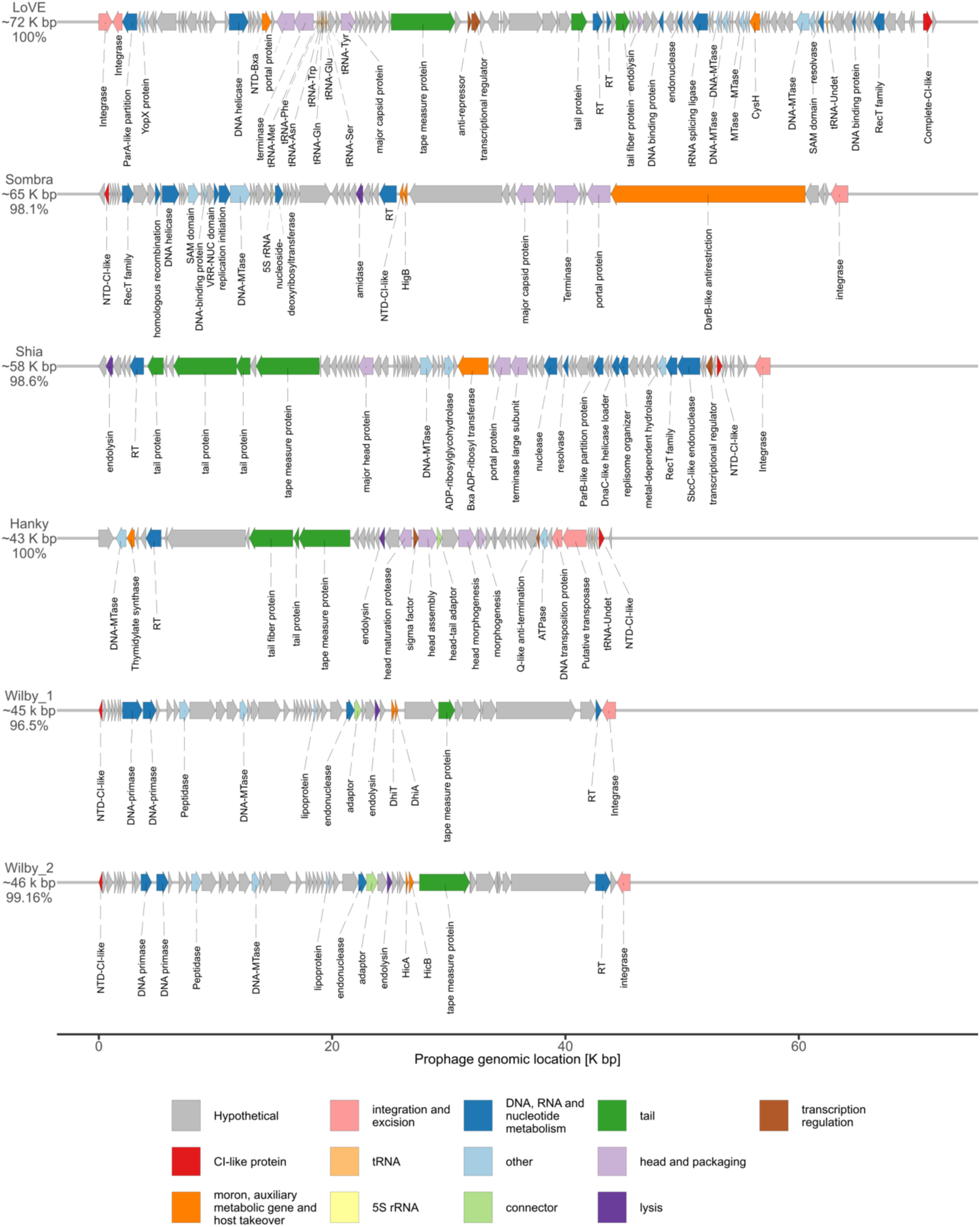
Bt_806 prophages. Schematic representation of the genomes of each of the six Bt_806 co-resident prophages. The size of each prophage genome and its CheckV completeness are written in this order below the phage VC. MTase: Methyltransferase, RT: Reverse transcriptase.

**Supplementary Figure 3.**
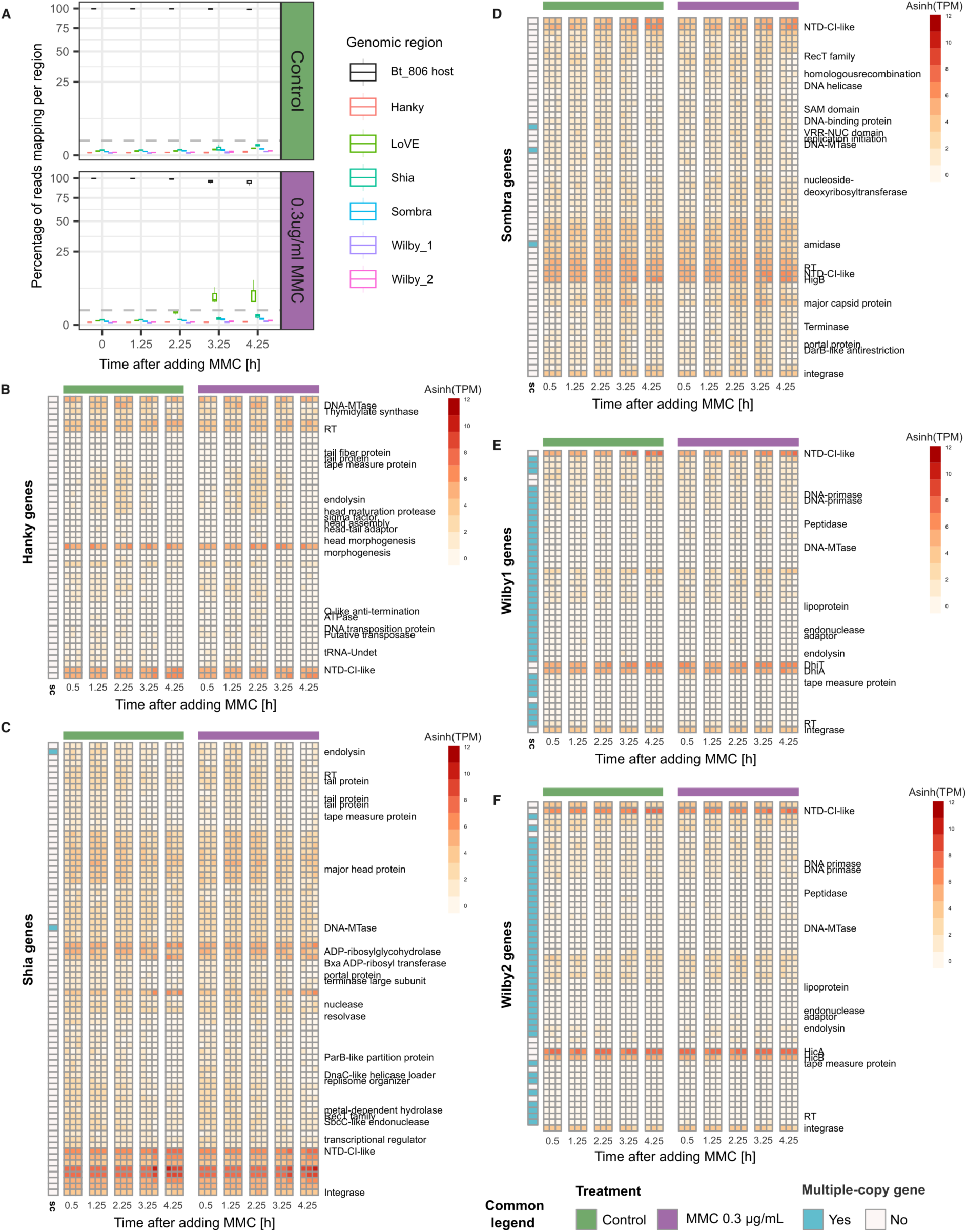
Only the LoVE prophage was induced upon DNA damage. **A)** Percentage of reads mapping to Bt_806 prophages and the genome. The dashed line represents 1% of the reads. Inverse hyperbolic sine of transcripts per million (TPM) of **B)** Hanky phage, **C)** Shia phage, **D)** Sombra, **E)** Wilby 1, and **F)** Wilby 2 in control (green bar on top) and MMC-treated (purple bar on top) samples across time points. Data from each biological replicate (n=3) is presented in separate columns at each time point. None of the genes were significantly different between treatments. The column at the left of the heatmap highlights genes that have multiple copies across the Bt_806 genome. Functional annotation is presented at the right of this last column; hypothetical/unknown genes were left blank.

**Supplementary Figure 4.**
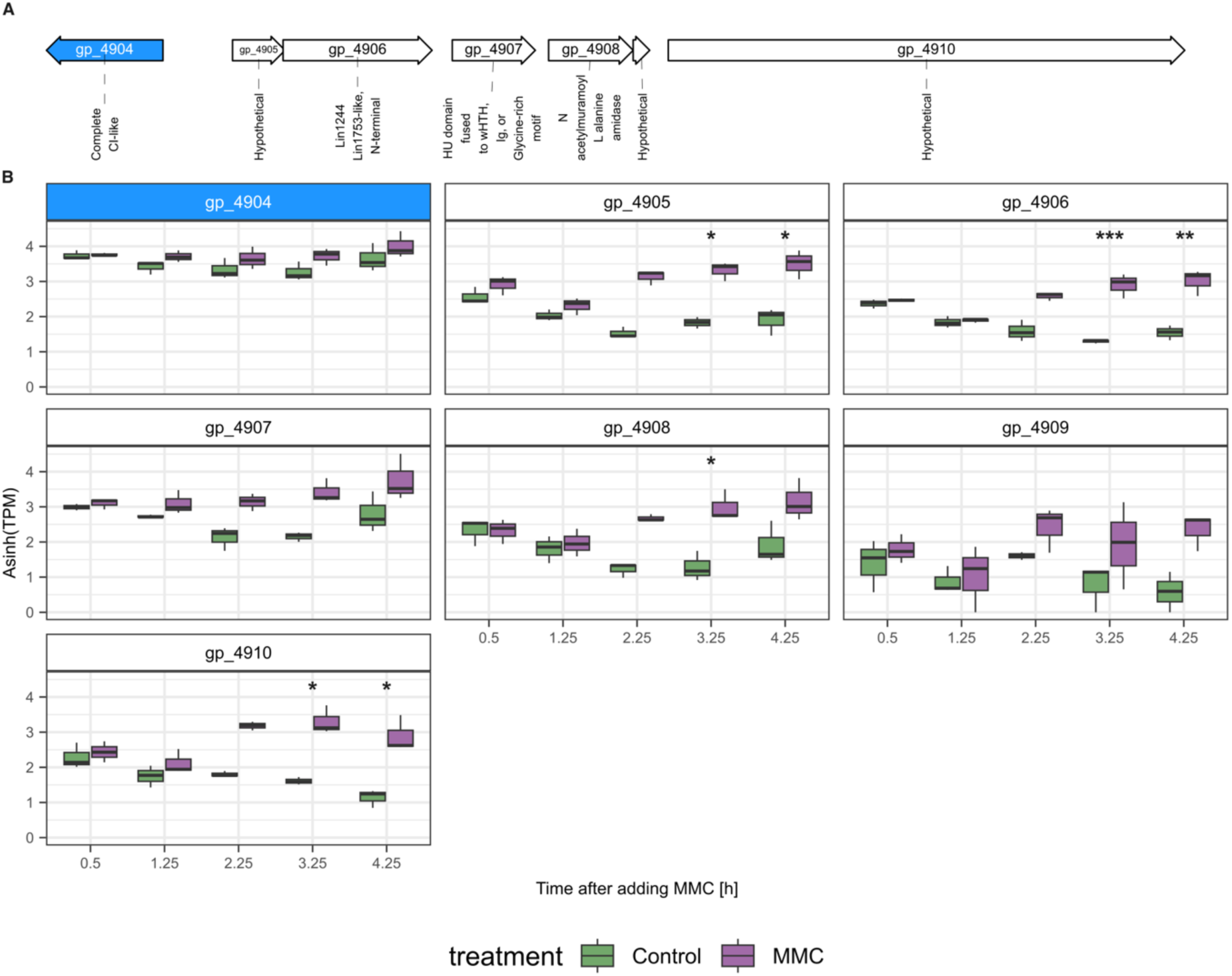
Bt_806 encodes a complete CI-like repressor in its genomic region that was not associated with a prophage. **A)** Schematic description of genes adjacent to the host-associated complete CI-like repressor. **B)** Expression levels of this operon during the experiment. Asterisks indicate the time points and genes whose expression was significantly higher in MMC-treated than control samples. *: FDR<0.05; **: FDR< 0.005; ***: FDR<0.0005, a quasi-likelihood (QL) negative binomial generalized log-linear model was fitted to the read-count data and QL F-test was employed to identified differential expressed genes. The complete CI-like repressor is highlighted in light blue in panel A and B.

**Supplementary Figure 5.**
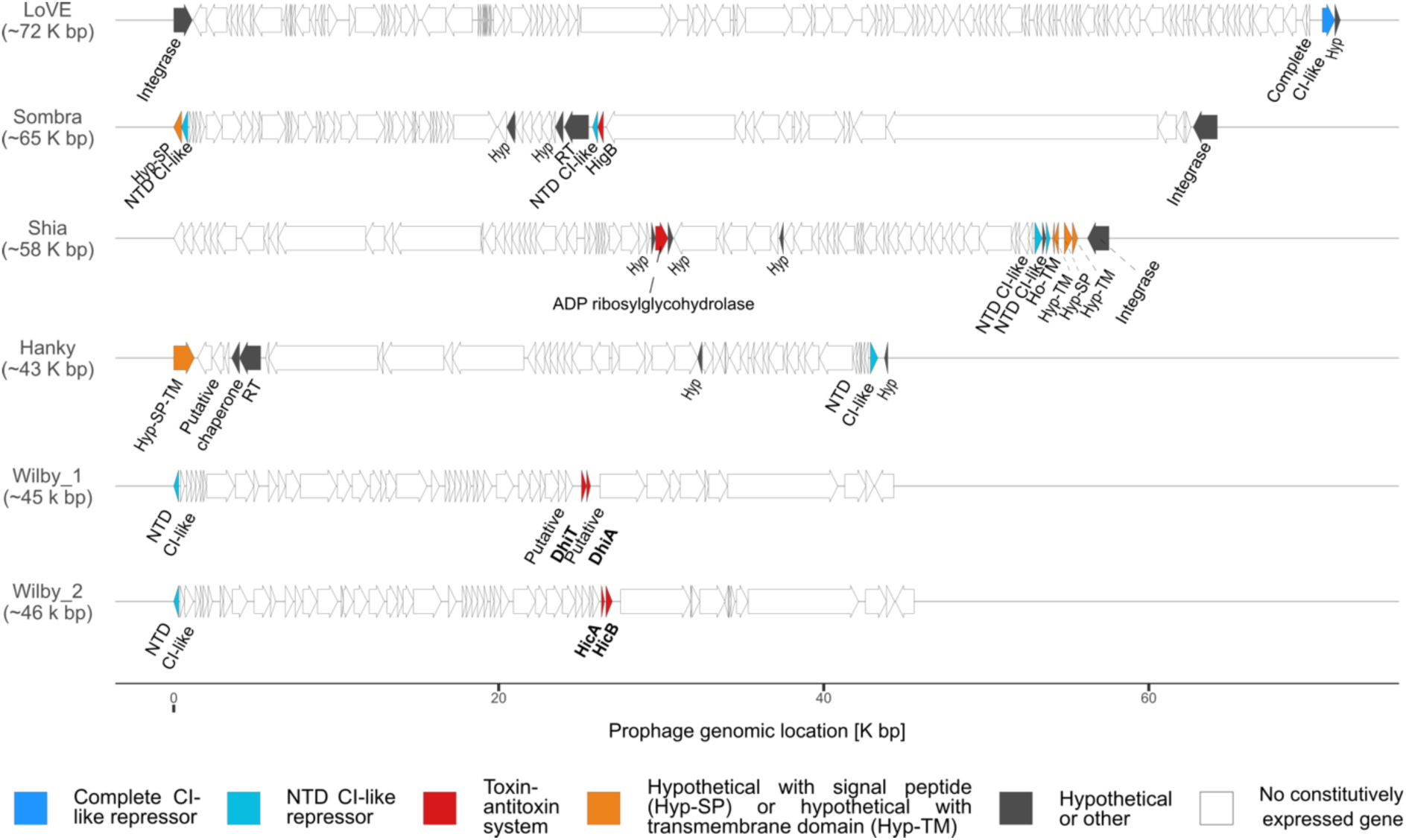
Transcriptionally active prophage-encoded genes during lysogeny. Constitutively expressed genes in Bt_806 prophages during stable lysogeny are coloured based on their annotation: Complete CI-like, in blue; incomplete CI-like, only containing NTD, in light blue; toxin-antitoxin systems (TA), in red; hypothetical protein (Hyp) with transmembrane domains (TM) or signal peptide (SP), in orange; and other, in grey. RT: reverse transcriptase. Wilby TA are in bold

**Supplementary Figure 6.**
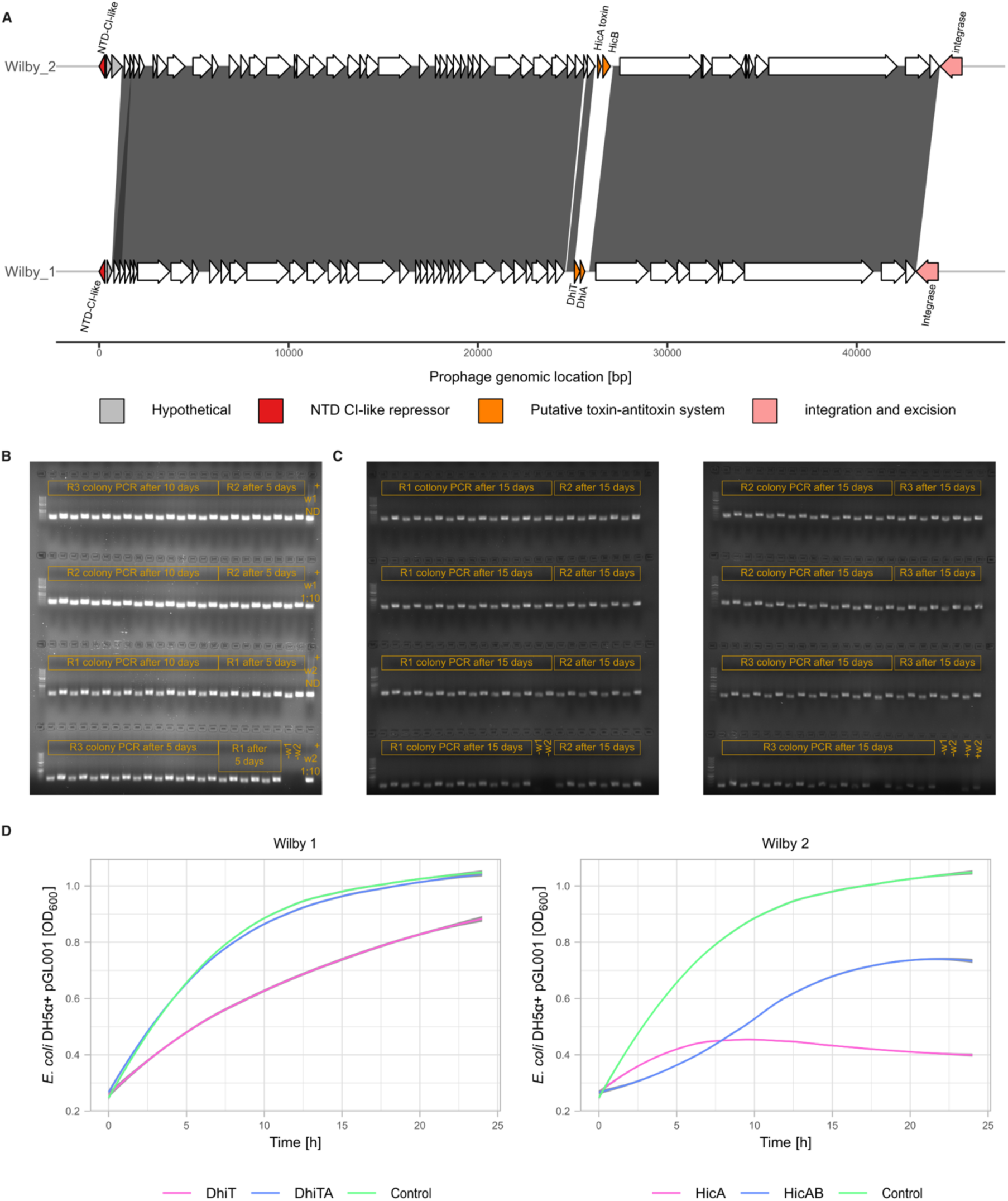
Comparison between the two Wilby prophages that co-reside in Bt_806 isolates. **A)** Grey shades indicate genomic regions that share more than 98% of nucleotide identity. Coloured and annotated genes are unique between the two Wilby prophages. Both Wilby prophages encode integrases that contain tyrosine recombinase domains, yet no significant amino acid similarity was found between them. **B-C)** Electrophoresis gels of Wilby colony-PCRs to check stability of both Wilby prophages through daily passages. Eight random colonies were sampled after 5 and 10 days of daily passages from each replicate (R1, R2, and R3) (n=24 per sampled day) (panel **B**), and 24 of them were sampled after 15 days (n=72) (panel **C**). Amplicon of each colony was loaded next to each other; the first amplicon corresponds to Wilby 1 and the second one to Wilby 2. Negative (-) and positive (+) controls were loaded as reference. ND=Input DNA was not diluted; 1:10 = Input DNA was 1:10 diluted. **D)** LOESS regression model for growth curves of *E. coli* DH5α cells expressing the putative toxin (pink), the toxin-antitoxin system (blue) of the two Wilby phages, or an empty pGL001 plasmid (green) as a control. All media contained chloramphenicol and aTc. Regression curves were calculated from four biological replicates. The solid line represents a locally weighted scatterplot smoothing (LOESS) curve with a span of 0.75. The shaded grey area indicates the 95% confidence interval.

**Supplementary Figure 7.**
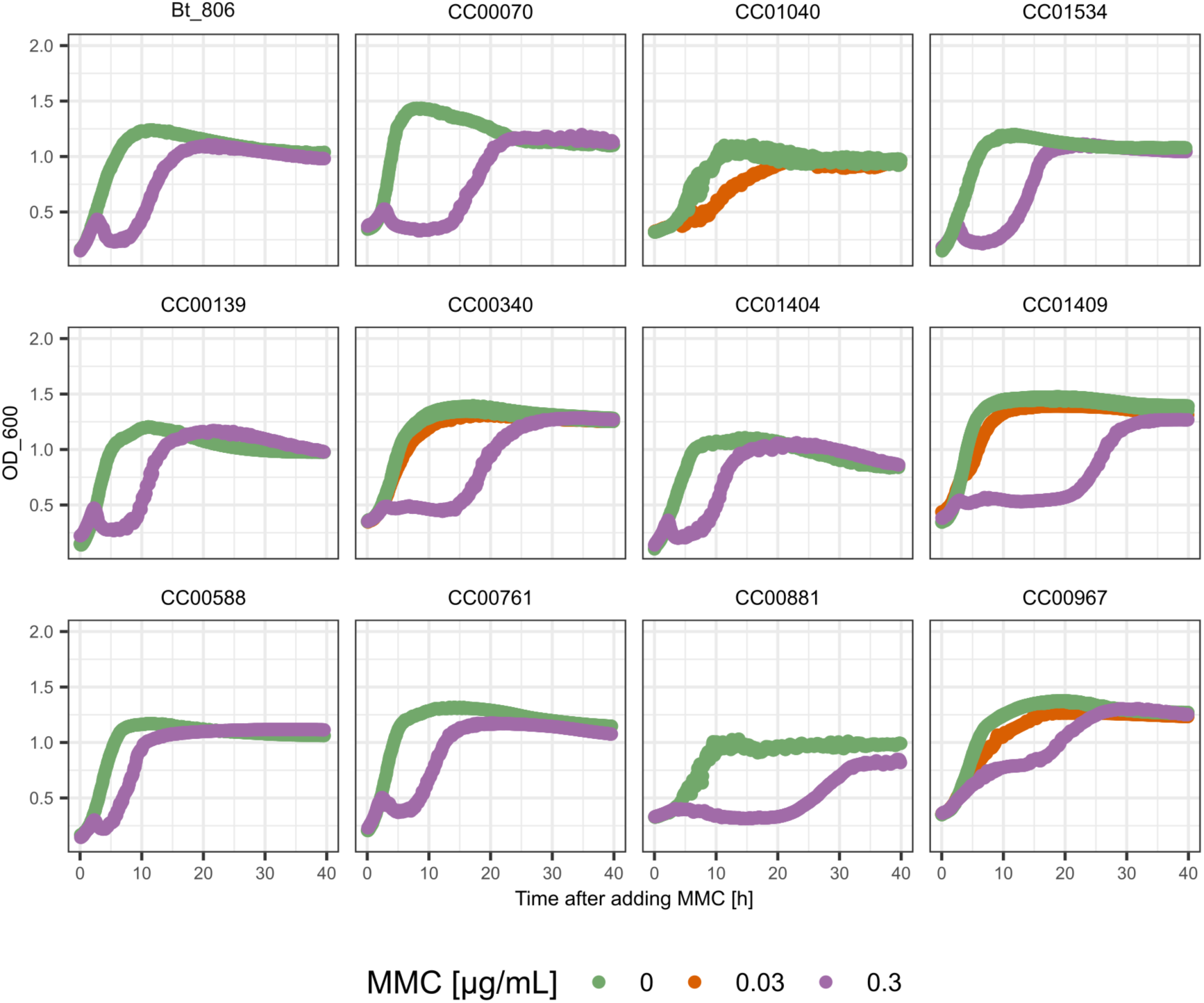
MMC impairs the growth of *Bacteroidales* poly-lysogens which carry only one phage encoding a complete CI-like repressor. Bacterial growth curves upon the addition of 0.03 ug/ml MMC (orange), 0.3 ug/ml MMC (purple), and control (green).

